# Data mining and classification of polycystic ovaries in pelvic ultrasound reports

**DOI:** 10.1101/254870

**Authors:** J. Jojo Cheng, Shruthi Mahalingaiah

## Abstract

**Objectives:** To develop and evaluate the performance of a rules-based classifier and a gradient boosted tree model for automatic feature extraction and classification of polycystic ovary morphology (PCOM) in pelvic ultrasounds

**Methods:** Pelvic ultrasound reports from patients at Boston Medical Center between October 1, 2003 and December 12, 2016 were included for analysis, which resulted in 39,093 ultrasound reports from 25,535 unique women. Following the 2003 Rotterdam Consensus Criteria for polycystic ovary syndrome, 2000 randomly selected ultrasounds were manually labeled for PCOM status as present, absent, or unidentifiable. Half of the labeled data was used as a training set, and the other half was used as a test set.

**Results:** On the test set of 1000 random US reports, the accuracy of rules-based classifier (RBC) was 97.6% (95% CI: 96.5%, 98.5%) and 96.1% (94.7%, 97.2%) for the gradient boosted tree model (GBT). Both models were more adept at identifying non-PCOM ultrasounds than either unidentifiable or PCOM ultrasounds. The two classifiers estimated prevalence of PCOM within our population’s ultrasounds to be about 44%, unidentifiable 32%, and PCOM 24%.

**Conclusions:** Although accuracy measured on the test set and inter-rater agreement between the two classifiers (Cohen’s Kappa = 0.988) was high, a major limitation of our approach is that it uses the ultrasound report text as a proxy and does not directly count follicles from the ultrasound images themselves.

## Introduction

Polycystic ovary syndrome (PCOS) has a population prevalence of 5-15%, depending on the diagnostic criteria used.^1^ It is characterized by hyperandrogenism, oligo-anovulation, and polycystic ovary morphology (PCOM). The 2003 Rotterdam Criteria define PCOM as the “presence of 12 or more follicles in each ovary measuring 2-9 mm in diameter, and/or increased ovarian volume (10 mL).”^2^ The exact follicle count cutoffs have since come into debate, as there is evidence that improved resolution in newer ultrasound technology increases the number of observable follicles, thus artificially increasing the prevalence of PCOM ^3^. Furthermore, PCOM is a relatively common finding in women that are healthy with a robust early follicular recruitment.^4, 5^

For these reasons, some authors have proposed changing the cutoff from 12 follicles to 20 or abandoning ultrasound altogether in favor of other biomarkers, such as serum anti-Mullerian hormone (AMH).^6, 7^ The ovarian volume cutoff of 10 ml may warrant revision as well, as it was based on 77 total control patients in the van Santbrink et al. and Pache et al. studies.

Existing electronic medical record (EMR) data may provide an opportunity for a closer study of the role of PCOM in the disease and suggest appropriate diagnostic standards. Compared to the high costs associated with collecting and interpreting new ultrasound data, hospital data is readily available, relatively inexpensive, and it can be linked with patients’ other clinical data for research into risk-reduction strategies. The major challenge with using EMR data in studying PCOS is that information about PCOM is captured in unstructured formats, such as ultrasound images and radiology text reports. Although “polycystic ovary syndrome,” the overarching disorder, appears in standard electronic health record ontologies (e.g. ICD-9 256.4 and ICD-10 E28.2), “polycystic ovary morphology,” the morphology specific to the ovaries, does not. Even if a radiologist may not be actively looking for PCOM and record its presence, its presence or absence can be inferred through reported volume measurements and descriptions of the internal structure. In this study, we develop and evaluate methods for automatic identification of polycystic ovarian morphology from pelvic ultrasound reports.

## Methods

### Data and study design

All ultrasounds from October 1, 2003 to December 7, 2016 were queried from the Boston Medical Center Clinical Data Warehouse. The start-date was selected to reflect the first day that ICD-9 codes were used and recorded at Boston Medical Center. This query yielded 39,093 ultrasounds for 25,535 unique patients; in total, this corpus consisted of 3,707,837 words, with each ultrasound containing an average of 95 words. A large majority of ultrasounds included 3-dimensional measurements of ovarian size and details about ovarian morphology, including the counts, echogenicity, and sizes of follicles and other ovarian abnormalities. Using a regular expression search, we found that out of 39,093 ultrasounds, only 6273 did not include three-dimensional volume measurements of the ovaries. These ultrasounds were still included for classification. The study was approved by the Institutional Review Board of Boston University School of Medicine and the Boston University Medical Campus.

### Reference label definition

Of the 39,093 ultrasounds, 2000 were randomly selected to be hand-labeled by a trained graduate-level researcher that was supervised by a board-certified Reproductive Endocrinologist and Infertility specialist. Of these, 1000 randomly selected ultrasounds were chosen to be a training set and another 1000 randomly selected ultrasounds were chosen to be a test set.

Following the Rotterdam criteria, an ovary was labeled with one of three determinations: as having polycystic morphology (PCOM), free of polycystic morphology (non-PCOM), or unidentifiable in the current examination (unidentifiable). We classified an ovary as PCOM if it was greater than 10 ml in volume in the absence of a dominant follicle (or other confounding structure), or if it contained a mention of classical PCOM appearance, such as “string of pearls orientation,” or “numerous peripheral follicles.” A determination of unidentifiable was made if the volume was greater than 10 ml, but there was also mention of a structure that confounded stromal volume (e.g. dominant follicle, dermoid cyst, hemorrhagic cyst, etc.), if there was no mention of ovaries, or if they were not measured. This follows the Rotterdam consensus that “If there is evidence of a dominant follicle (>10mm) or a corpus luteum, the scan should be repeated the next cycle.” An ovary was considered non-PCOM if it was smaller than or equal to 10 ml and did not contain mention of classical PCOM appearance.

### Modeling

To accomplish the classification task, two approaches were used: (1) a rule-based classifier (RBC) that checks the size of each ovary and detects important words appearing close to the word, “ovary” which may indicate the presence of PCOM or confounding structures, and (2) a gradient boosted tree model (GBT) that uses a feature vector of term frequencies and each side’s ovarian volume. The overall architectures are depicted in Figures 1 and 2, respectively.

### Rule-based classifier (RBC)

The rule-based classifier consists of several modules: a preprocessing phase, contextualization, volume extraction, corpus and document-term matrix creation, term selection, detection, and final evaluation. The overall scheme is given in Figure 1.

### Text preprocessing

In the preprocessing phase, standard NLP techniques for cleaning and tokenization are used, including removing irregular characters, nonstandard spaces, punctuation, stopwords and line breaks. Each word is then stemmed with Porter’s Snowball stemmer. This phase ensures that these irregularities do not affect the analysis. An overview of standard preprocessing techniques is given in Manning et al., 2008.^8^

### Contextualization, volume extraction, corpus creation

In the contextualization module, in order to limit the number scope of text used for prediction, only the sentences mentioning the word “ovary,” as well as the sentences immediately succeeding such sentences are extracted. This is justified because in a pelvic ultrasound report, descriptions of other organs such as the uterus or kidneys are not relevant for inferring polycystic ovarian morphology. Next, the program uses regular expressions to assign sentences and phrases to the left and right ovary for analyzing the two separately. The product of this module is a data frame, with up to two rows of the data frame corresponding to an ovary mentioned in each ultrasound. Sentences that describe both ovaries appear once in each ovary.

During the volume extraction phase, regular expressions are used to extract 3-dimensional measurements from the ultrasounds and to determine the correct unit of measurement. Volume of ovaries are calculated according to the formula, *length · width · height* · 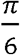, as recommended for calculating the volume of an (ellipsoid) ovary.^9^ Ovarian volume measurements with only 1- and 2-dimensions reported are excluded. The corpus is generated such that the group of sentences about each unilateral ovary corresponds to each document in the corpus. Thus, each ultrasound report generated at most two documents in the corpus.

### Document-term-matrix (DTM) creation

In order to make use of the volume cutoff, we looked to eliminate cases of ovaries enlarged by other pathologies, such as hemorrhagic cysts and dermoids. For accurately distinguishing between ovaries enlarged by increased stromal volume versus ovaries enlarged due to recruitment of a dominant follicle or presence of abnormal pathology, we needed a comprehensive list of reasons that could confound the representation of stromal volume through total ovarian volume.

This list was constructed empirically by creating a document-term matrix, and finding 2-, and 3-grams (two word phrases, and three word phrases) correlated with a large ovarian volume.^10^ We chose not to use 1-grams in order to decrease statistical variance; for example, the presence of the word “cyst” does not indicate whether the actual presence or absence of a cyst is being described in the text. After sorting into large (>10 ml) and small (<10 ml) ovaries, 2- and 3- grams appearing in at least 18 (0.1%) of big ultrasounds were extracted.

Many ultrasounds also included sentences with language similar to the following: “numerous follicles arranged in a peripheral distribution indicative of PCOS.” The list of 3-grams correlated to “numerous peripheral follicles” and the list of 2-grams most correlated with the phrases, “ovarian syndrome,” “polycystic ovarian,” and “string of pearls” were extracted in order to detect the noted presence of PCOM.

This approach offers the distinct advantage of generating all possible reasons appearing in the dataset, and in this way implicitly accounting for the natural language of the data. Typically this problem would be impossible to solve *a priori*.

### Term selection and detection

The previous step reduced the corpus to essentially a list of a few thousand n-grams. This list of n-grams was then curated so only phrases that indicate the presence of polycystic morphology or the presence of a volume confounder were kept (Supplemental Tables 1 & 2). During the detection phase, each of these phrases then counted as a vote for the presence of volume confounding or polycystic morphology. This selection of n-grams was iteratively refined on the training set. The rules for the detection algorithm followed the Rotterdam Consensus Criteria and used the ovarian volume and presence of numerous follicles to determine the presence of PCOM for each ovary (Supplemental Figure 3). Overall patient status for the given ultrasound is determined from the status of both ovaries. The patient is determined to have PCOM on a given ultrasound if at least one ovary had PCOM.

### Evaluation

During the evaluation phase, predicted classification labels were compared to the hand-labeled test set of 1000 ultrasounds for computing confusion matrices and accuracy statistics.

### Gradient boosted tree classifier (GBT)

Gradient boosting is a machine learning technique that ensembles many weak prediction models – in this case, decision trees – in an additive, stage-wise manner to produce a stronger model. At each stage, new weak models are introduced to account for the errors of previously existing weak models.^11, 12^ The schematic for gradient boosted tree model is given in Figure 2. The GBT uses the same preprocessing module as the RBC. After the ultrasound text is preprocessed, the corpus is generated without the contextualization and splitting into left and right ovaries: each ultrasound report corresponds to one document within the corpus. This was done to see if the bilateral nature of detecting PCOM could be modeled by the algorithm without explicitly programming it. Similarly, a document-term matrix was generated from all 1-, 2-, and 3-grams without any human selection to see if important phrases could be learned during training time.

The contextualization and volume extraction modules were used to extract ovarian volumes as predictors for the boosted tree algorithm. The GBT used the XGBoost library with default parameters except for the the number of training iterations, which was selected by minimization of 5-fold cross-validation (CV) error on the training data.^13^ Hyperparameter tuning was briefly attempted, but quickly abandoned because there were no significant gains in CV training error compared to the extra training time required.

## Results

Table 1 includes the summary information about the source population at Boston Medical Center which contributed ultrasounds, including demographics and general health. Compared to other cohorts, this group has significant race/ethnic diversity, with nearly 80% non-white participants, and significant socio-economic diversity, with a third of the patients not finishing high school, another third attaining high school diplomas or a GED, and about a third with at least some higher education. Nearly 2% have been homeless, compared to 0.5% of the United States population.^14^ The diversity of the cohort reflects the diverse patient population served by Boston Medical Center, which serves as the region’s safety net hospital. For markers of general health, blood pressure is within a normal range. About a third of the patients are underweight or normal, another third are overweight, and a third are obese.

**Table 1:** Source population characteristics.

| N = 25,535 | % (n) or mean (SD) |
| --- | --- |
| **Age** |  |
| Age at ultrasound | 31.4 (7.9) |
| **Race** |  |
| Asian | 3.5% (883) |
| Black/African American | 48.9% (12,489) |
| Hispanic/Latino | 19.9% (5076) |
| Non-Hispanic White | 21.0% (5358) |
| Other*/Multiracial | 5.6% (1432) |
| Declined/Not Available | 1.2% (297) |
| **Vitals** |  |
| BMI (n = 18443) |  |
| Underweight < 18.5 | 1.2% (225) |
| Normal weight 18.5-24.9 | 27.9% (5138) |
| Overweight 25-29.9 | 30.4% (5610) |
| Obese Class I 30-35 | 20.7% (3818) |
| Obese Class II 35-40 | 11.2% (2060) |
| Obese Class III >40 | 8.6% (1592) |
| SBP (mmHg) (n = 21353) | 118.8 (11.6) |
| DBP (mmHg) (n = 21351) | 75.2 (7.6) |
| Age at Menarche (n = 178) | 12.7 (2.0) |
| **Socioeconomic Status** |  |
| Education (n = 39,638) |  |
| Did not attend school | 5.6% (1416) |
| 8<sup>th</sup> grade or less | 4.6% (1169) |
| Some high school | 22.3% (5690) |
| Graduated high school or GED | 27.1% (6928) |
| Some college/vocational/technical school | 13.4% (3426) |
| Graduated college/postgrad. | 16.8% (4292) |
| Other Education | 1.0% (266) |
| Declined/Not Available | 9.2% (2348) |
| Ever homeless |  |
| Yes | 1.9% (486) |
| No | 98.1% (25,040) |
| Not available | <0.1% (9) |

\*Includes groups with less than 250 members: American Indian/Native American, Middle Eastern, and Native Hawaiian/Pacific Islander
SBP = Systolic Blood Pressure; DBP = Diastolic Blood Pressure

Table 2 presents the performance of the two models on the dataset. The overall test set accuracy of the RBC was 97.6% (95% CI: 96.5%, 98.5%) and 96.1% (94.7%, 97.2%) for the GBT. The RBC outperforms the gradient boosted trees in correctly classifying ultrasounds as unidentifiable and PCOM. Both models were more adept at identifying non-PCOM ultrasounds than either unidentifiable or PCOM ultrasounds. Table 3 presents classification frequencies predicted by the two classifiers for the entire dataset and their confusion matrix. Inter-rater agreement between the two classifiers was high (Cohen’s Kappa = 0.988) and the two classifiers estimated prevalence of PCOM within our population’s ultrasounds to be about 44%, unidentifiable 32%, and PCOM 24%.

**Table 2:** Classification metrics across different models.

| Model | Accuracy | Class label | Sensitivity | Specificity | PPV | NPV |
| --- | --- | --- | --- | --- | --- | --- |
| Gradient boosted trees | 0.961<br>(0.9471, 0.9721) | No PCOM | 0.9977 | 0.9982 | 0.9977 | 0.9982 |
|  |  | Unidentifiable | 0.9474 | 0.9705 | 0.9387 | 0.9748 |
|  |  | PCOM | 0.9146 | 0.9761 | 0.9259 | 0.9723 |
| Rules-based classifier | 0.976<br>(0.9645, 0.9846) | No PCOM | 0.9885 | 0.9982 | 0.9977 | 0.9912 |
|  |  | Unidentifiable | 0.966 | 0.9852 | 0.9690 | 0.9838 |
|  |  | PCOM | 0.9668 | 0.9829 | 0.9472 | 0.9894 |

PPV = Positive predictive value; NPV = Negative predictive value

**Table 3:**
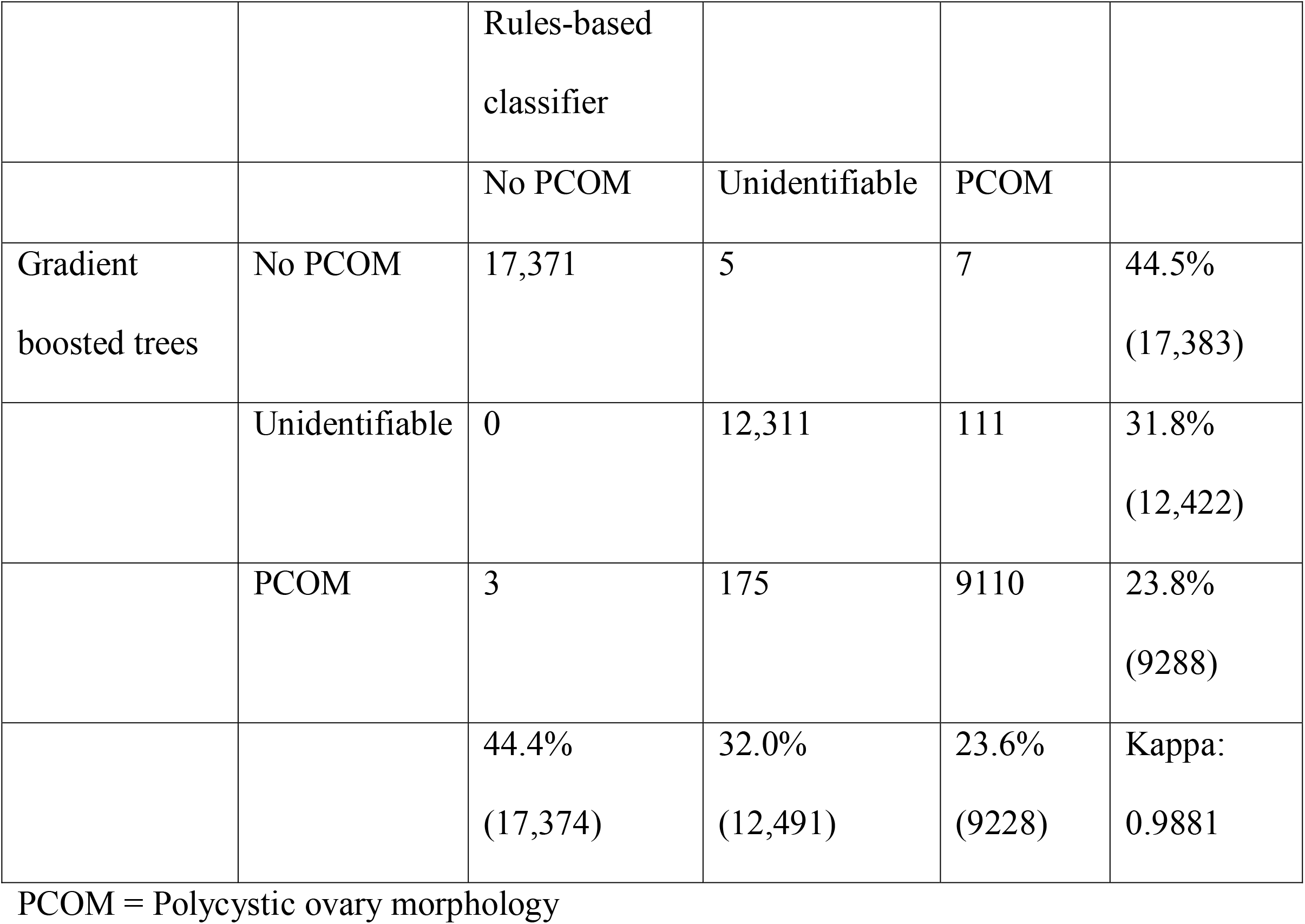
Classification frequencies by model and confusion matrix.

## Discussion

The two methods for detecting the presence of polycystic ovarian morphology in ultrasound reports demonstrated good sensitivity and specificity on our test set. The rules-based classifier performed slightly better than the gradient-boosted trees (97.6% vs 94.7%), but a weakness of the method is that it takes longer to develop. Both methods likely work well within the current application because the highly sequential nature of the Rotterdam Consensus Criteria for identifying PCOM is amenable to decision trees.

The RBC was likely successful because the corpus could be reduced to a relatively small set of 600 n-grams (Supplemental Tables 1 & 2). Thus, the problem was reduced to that of marking those that suggest the presence of volume confounders or the presence of small, peripherally oriented follicles. This approach may generalize well to other applications where the central problem is in identification of diseases that contain specific language for the condition in question.

Upon closer inspection of the RBC errors, it appears that the algorithm has difficulty with temporal language. For example, it may recognize that a radiologist is writing about a large cyst, but it will not recognize that this large cyst was present only in the past and is no longer seen in the current exam. It also has considerable difficulty with classifying ultrasounds when the ovarian volume is not reliably extracted, either due to program errors from not recognizing uncommon phrasing or typographical error very difficult for a program to parse. These typographical errors seemed to have arisen from either mistakes in typing or dictation software. For example, “ovary measures 4.3×1.6x<u>approximately</u> 3 cm” or “left ovary appears…measuring 2.7×1.7 <u>a</u> 2.1 cm” or “the right ovary measures 2<u>.xby</u> 2.8×1.6 cm” (errors underlined).

Since it relied on the same volume extraction protocol, the GBT model also struggled when the volumes were not properly extracted. Both models were surprisingly error tolerant, because often classifying a single ovary correctly was enough to classify the overall status. This aspect of classifying PCOS is probably responsible for the asymmetrical importance rating given by the GBT to each ovarian volume (Supplemental Figure 4). Other errors of the GBT model seemed to arise from the limitations of a bag-of-words method. Since only 1-, 2-, and 3-grams were used as predictors, it had difficulty distinguishing when description of a cyst appeared in the left or right ovary without being able to detect long-range word interactions. For example, the text for one ultrasound reads, “the right ovary…containing 3.5×4 cm hemorrhagic cyst. the left ovary measures 3×3×5 cm.” The RBC would have split these two sentences and assigned them to each ovary, but the GBT simply considers the entire set of n-grams, which would capture the three words, “cyst. left ovary” as a single unit, ultimately guessing that a cyst is present in the left ovary. Supplemental Figure 4 shows the most important predictors learned by the GBT model.

When predictions for the rest of the full dataset were generated by the two methods, the two classifiers agree strongly on what does not count as PCOM. Most of the disagreement is on a subset that of 286 cases that seem to be either PCOM or unidentifiable. Out of the 1000 ultrasounds in the test set, there were 16 cases that were misclassified by both.

We suspect that the test performance of the RBC model is slightly misleading: the test set accuracy is higher than the accuracy on the training set used to refine the RBC. We expect actual performance of the RBC model to be closer to the accuracy measured on the combined training and test set, which is 0.968. The overly optimistic assessment of performance seems to stem from the test set containing examples that were “easier” for the RBC to classify due to random chance.

We reiterate that having PCOM does not imply that a patient has PCOS and that the women in the approximately 9200 ultrasounds labeled with PCOM do not necessarily have PCOS. The 24% prevalence of PCOM in our dataset is comparable to the 22% PCOM prevalence reported by Clayton et al. in unselected women,^4^ and 23% reported by Polson et al. in women who considered themselves healthy.^15^ As our dataset is a hospital-based sample, the women in our dataset likely had other preexisting conditions that led them to seek care, which biases our sample towards poorer health.

Furthermore, inferring PCOM status from ultrasound reports and not the ultrasounds themselves can be inexact, because a radiologist may not explicitly describe everything that appears in the image, only aspects of the ultrasound relevant to the indication. Thus, the absence of an object from the ultrasound report does not guarantee its absence in the image. Another limitation is that the algorithms are unable to explicitly count follicles (since explicit follicle counts are not usually given), and instead uses certain language about “numerous follicles” as a proxy. Ascertainment of follicle counts is critical to more sensitive and specific automatic identification of PCOM. These algorithms only detect morphology, and using them for diagnosing PCOS would also require the exclusion of those who are using hormonal contraceptives, currently pregnant, or diagnosed with another endocrinopathy.^1, 16^

Current open questions about PCOM include characterizing the within-woman variability of ovarian volume measurements over time, understanding race-ethnic characteristics of follicle counts, the association between polycystic ovaries with metabolic disease, and the relationship between ovarian phenotype with polycystic ovary syndrome. Historical ultrasound data collected as part of natural clinical processes could be a source of answers to these inquiries, but methods for harnessing this data are lacking. Here we developed two classifiers based on NLP techniques that predict PCOM state with great sensitivity and specificity. In particular, our classifiers provide a means of answering the first question and investigating the others. We recommend that further research in automatic feature extraction for PCOS be directed towards methods for the raw ultrasound images themselves.

## Acknowledgements

We would like to acknowledge Linda Rosen, the research manager of the Clinical Data Warehouse for procuring the dataset.

---

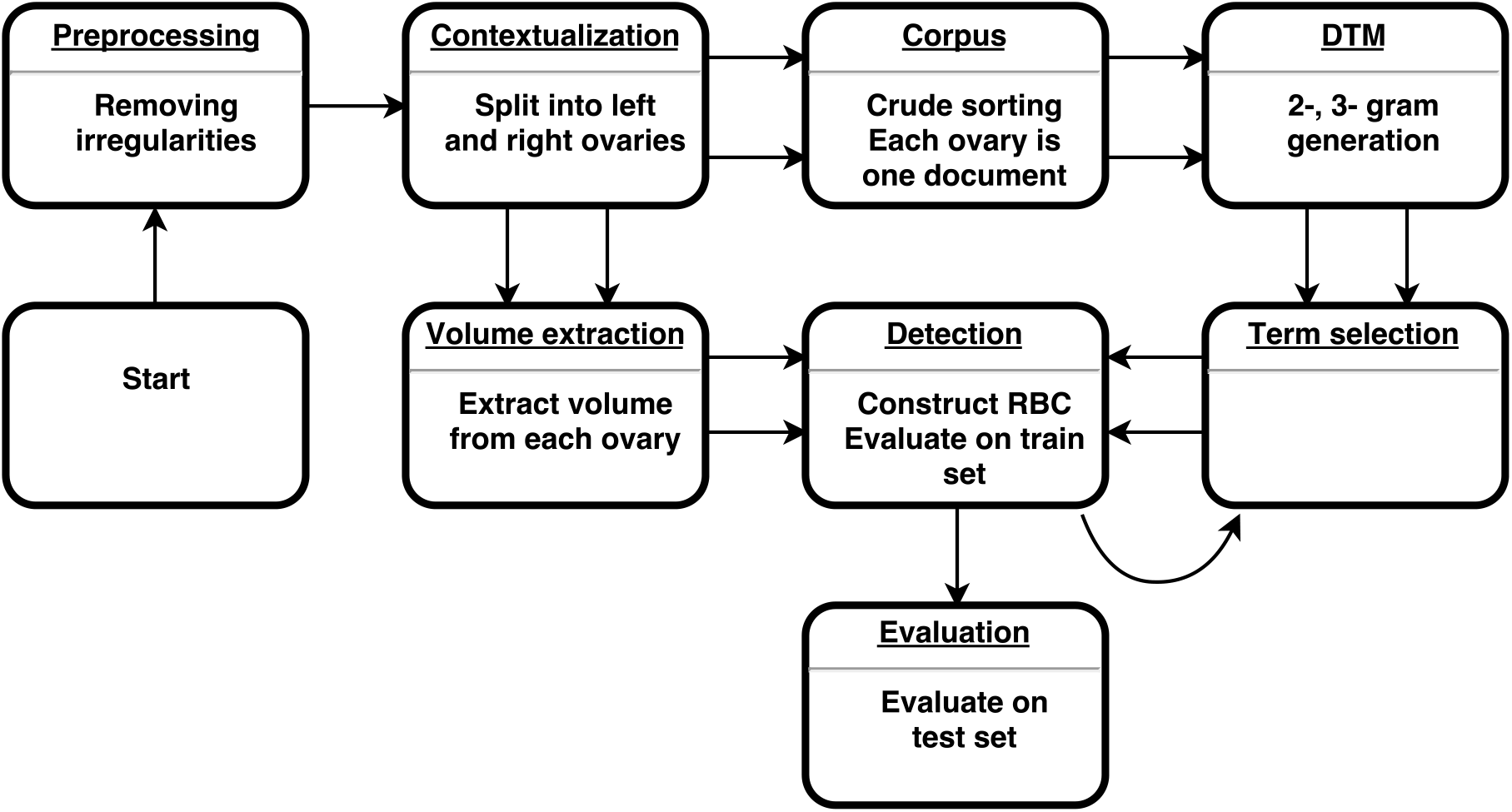

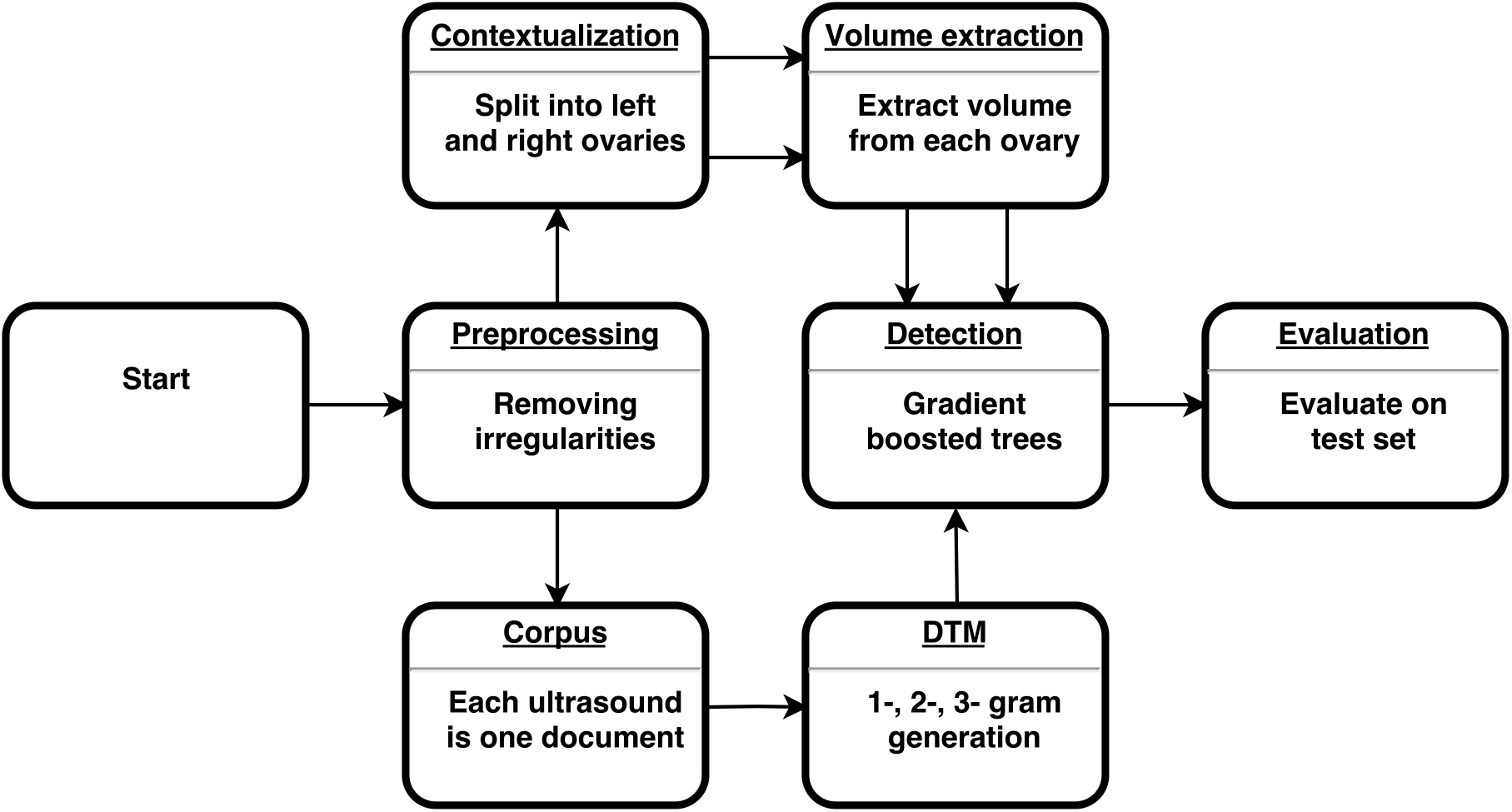

